# Preliminary analysis of the metabolic and physical activity profiles of mice lacking the *slc43a3*-encoded equilibrative nucleobase transporter 1

**DOI:** 10.1101/2025.04.17.649357

**Authors:** Aaron L. Sayler, James R. Hammond

**Affiliations:** Department of Pharmacology, Faculty of Medicine & Dentistry, University of Alberta, Edmonton, Alberta, Canada

## Abstract

*SLC43A3* encodes for a membrane transporter selective for purine nucleobases (equilibrative nucleoside transporter 1; ENBT1). Adenine, an endogenous substrate for ENBT1, plays an important role in many biochemical and physiological processes, including cellular energy metabolism. To investigate how the loss of ENBT1 impacts these processes, we generated a *slc43a3*-null (global; KO) mouse model.

Metabolic function, physical activity, and food and water consumption were assessed in male and female wild-type (WT) and KO mice (age 10-12 weeks) for a 66-hour period (12 hr light/dark cycle). Blood pressure and heart rate of each group of mice were also assessed using a rodent tail cuff method.

Female KO mice showed a significant decrease in oxygen consumption and carbon dioxide generation with no change in RER, relative to female WT mice. This was accompanied by a significant 4-hour negative phase-shift in circadian rhythm in the metabolic and activity measures in the female, but not male, KO mice. Male KO mice, in contrast, displayed a significant decrease in rearing activity and blood pressure, and an increase in metabolic expenditure relative to WT mice.

It may be concluded that loss of *slc43a3*-encoded ENBT1 impacts numerous measures of activity in mice, with female mice impacted more significantly than male mice. This may reflect disruption of purinergic processes associated with energy metabolism coincident with changes in adenine availability. The results of this study also imply an interaction between estrogen and adenine metabolism in the regulation of circadian rhythms.

## Introduction

*SLC43A3* encodes for a membrane transporter selective for purine nucleobases (equilibrative nucleoside transporter 1; ENBT1)[1, 2]. Adenine, an endogenous substrate for ENBT1, plays an important role in many biochemical and physiological processes, including serving as a precursor for the intracellular production of adenine nucleotides[3, 4]. Adenine nucleotides are integral to cellular energy metabolism and adenosine has a host of biological regulatory functions including being involved in vaso-regulation and neuronal modulation[5]. Therefore, it is not unreasonable to expect that the genetic deletion of *slc43a3* in mice would have a significant impact on cellular purine availability and biological functions that are regulated by purine nucleosides and nucleotides. We have recently published on the creation and use of a *slc43a3*-null (global) mouse model to assess the role of the encoded transporter in the absorption of orally administered 6-mercaptopurine[6]. These *slc43a3*-null mice were found to be viable with no obvious defects in gross morphology. We now report on an analysis of the metabolic energy profile, food and water consumption, mobility, and cardiovascular impact of the loss of *slc43a3* in these mice.

## Materials and methods

*Slc43a3*-null (KO) and wild-type (WT) C57BL/6J mice were bred in-house via heterozygous pairings as described previously[6]. Food and water were provided ad libitum and standard housing was used. All animal work was conducted using protocols approved by the Canadian Council of Animal Care via the Faculty of Medicine and Dentistry, University of Alberta, Animal Use Committee.

A comprehensive lab animal monitoring system (CLAMS-HC, Columbus Instruments, Columbus, OH) was used in tandem with a Columbus Instruments Oxymax Lab Animal Monitoring System software (Cardiovascular Research Centre, University of Alberta) to measure total horizontal volitional movement, ambulatory activity (multiple beam breaks), Z-activity (rearing), food intake, water intake, volume of O_2_ consumption (VO_2_), volume of CO_2_ production (VCO_2_), respiratory exchange ratio (RER) and energy expenditure. Mice had access to standard chow and water and were singly housed at room temperature for these analyses. Eight mice of each sex and genotype (WT, KO) were assessed at 10-12 weeks of age. Mice were acclimated to the metabolic cages for 24 hours, and the above noted parameters were then measured continuously (every 1-3 minutes), with 12-hour light/dark cycles, over a 3-day period starting at 7:00 pm (beginning of dark cycle; designated as zeitgeber time zero) on the first day to 1:00 pm on the third day. For analysis, the data obtained was binned by hour (averaged for VO_2_, CO_2_, RER, and energy expenditure; summed for activity measurements and food and water consumption). Differences were assessed based on statistical analyses (one-way ANOVA with Tukey’s post test, P<0.05) of the area under the curve (AUC), determined using GraphPad Prism v10.4, and the circadian phase shift and fluctuation amplitude obtained from cosinor analysis using the FFT NLLS method[7] with linear detrending (biodare2.ed.ac.uk) [8]. The differences in baseline derived from the cosinor analysis also provided a measure of the average change in the measured parameter, similar to the AUC values. However, since the AUC measurement uses the actual data points, as opposed to an extrapolated value for derivation of the cosine baseline, the AUC was considered to be the more rigorous statistical comparison.

Cardiovascular parameters (systolic and diastolic blood pressure, heart rate) were measured using a CODA^®^ High Throughput (Kent Scientific, Torrington, USA) non-invasive blood pressure measurement system designed specifically for rodents, by the Cardiovascular Research Centre, University of Alberta. A total of 8 WT mice and 8 KO mice of each sex were analyzed at 10-12 weeks of age. Animals were placed in the CODA^®^ Animal Holders wearing the CODA^®^ High Throughput occlusion cuffs and volume-pressure recording cuffs on a heating pad (35°C) and covered with an opaque insulating sheet for calming effect to assist with attaining normal physiological blood flow. Cardiovascular parameters were recorded once a minute for a total of 15 minutes. All animals were acclimated in this facility for 15-minute periods on 3 successive days preceding the experiments to familiarize them with the procedure and reduce stress-induced cardiovascular anomalies.

## Results

The data obtained from these studies were assessed for both the magnitude of the change resulting from loss of *slc43a3*, and the effect of this gene deletion of circadian rhythm. Differences in the parameters between male and female mice were also evaluated. In terms of changes in magnitude (AUC analyses), female WT mice had a significantly higher (by approximately 13%) overall VO_2_ and VCO_2_ compared to male WT mice, but there was no significant difference between male and female KO mice in that regard (Fig 1, Table 1). This change was due to a significant 10% reduction in VO_2_ in the female KO mice relative to female WT mice, and a similar trend towards a reduced VCO_2_ in the female mice with the loss of *slc43a3* (Table 1). The female KO mice also showed an approximate 20% decline, relative to female WT mice, in the magnitude of the fluctuations (cosine amplitude) in both VO_2_ and VC0_2_ over the 24-hour cycle (Fig 1, Table 1). Due to the parallel changes in VO_2_ and VCO_2_ in these subject groups, there was no significant difference in respiratory exchange ratio (RER) (Table 1). Loss of *slc43a3* also led to a significant 13% increase in metabolic energy expenditure in the male mice, but not in the female mice (Table 1). This increase in energy expenditure was accompanied by a significant decrease in rearing activity and a trending decrease in activity in general in the male KO mice that was not seen for the female KO mice (Fig 2, Table 1). There was no real impact of the loss of *slc43a3* on feeding or drinking behaviour other than a slight decrease in cosine fluctuation amplitude in water consumption by the female KO mice, relative to female WT mice (Fig 3, Table 1).

**Fig 1.**
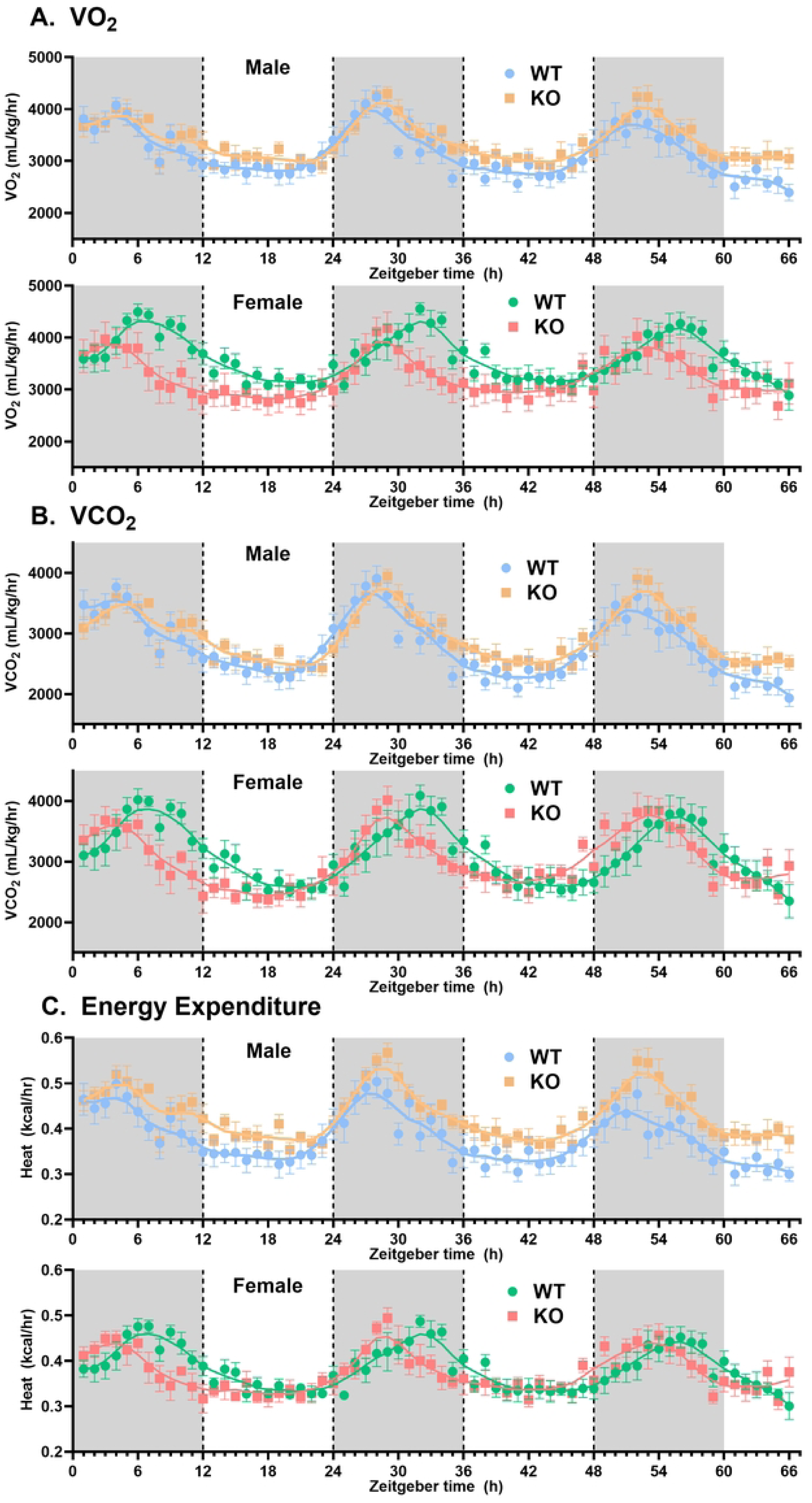
Effect of *slc43a3* deletion on metabolic performance parameters in male and female WT and KO mice. Using an automated Columbus Lab Animal Monitoring System (CLAMS), oxygen consumption (VO_2_) and carbon dioxide generation (VCO_2_) were measured over 66 h (12 h L/D) with zeitgeber time 0 defined as the start of the first dark phase. The values for respiratory exchange ratio (RER) and energy expenditure were derived from these data. Each point is the mean ± SEM from 8 mice. The lines represent LOWESS curve fits to the data using a 20-point smoothing window (GraphPad Prism v10.2). The area under the curves for these data sets are shown in Table 1.

**Fig 2.**
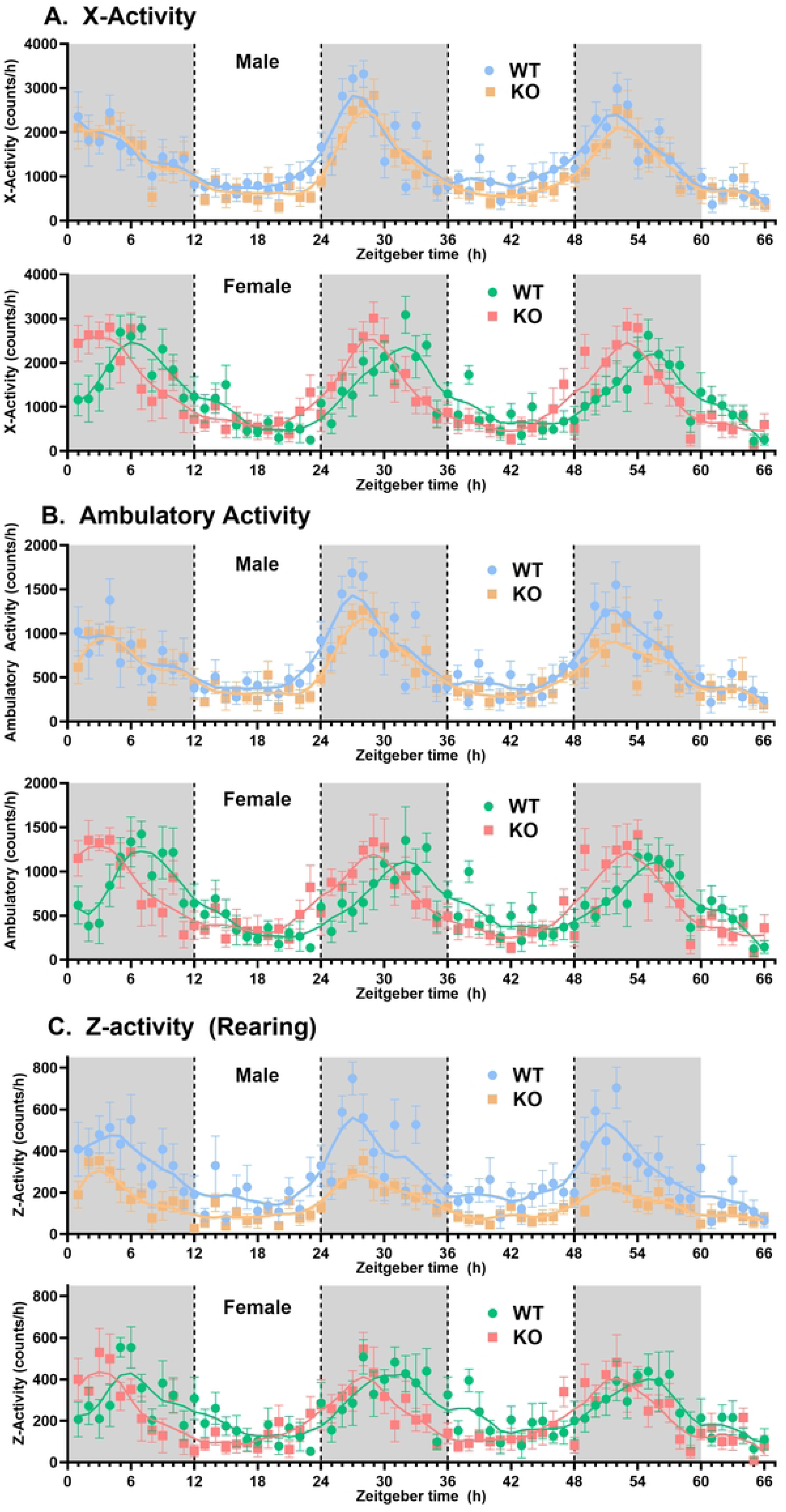
Effect of *slc43a3* deletion on physical activity of male and female WT and KO mice. Using an automated Columbus Lab Animal Monitoring System (CLAMS), the total number of horizontal beam breaks (X-Activity), the number of consecutive horizontal beam breaks (Ambulatory Activity), and the number of elevated beam breaks (Z-Activity, representing rearing activity of the mouse) were measured continuously over 66 h (12 h L/D) and binned to obtain the beam breaks (counts) per h. Zeitgeber time 0 was defined as the start of the first dark phase. Each point is the mean ± SEM from 8 mice. The lines represent LOWESS curve fits to the data using a 20-point smoothing window (GraphPad Prism v10.2). The area under the curve for these data sets are shown in Table 1.

**Fig 3.**
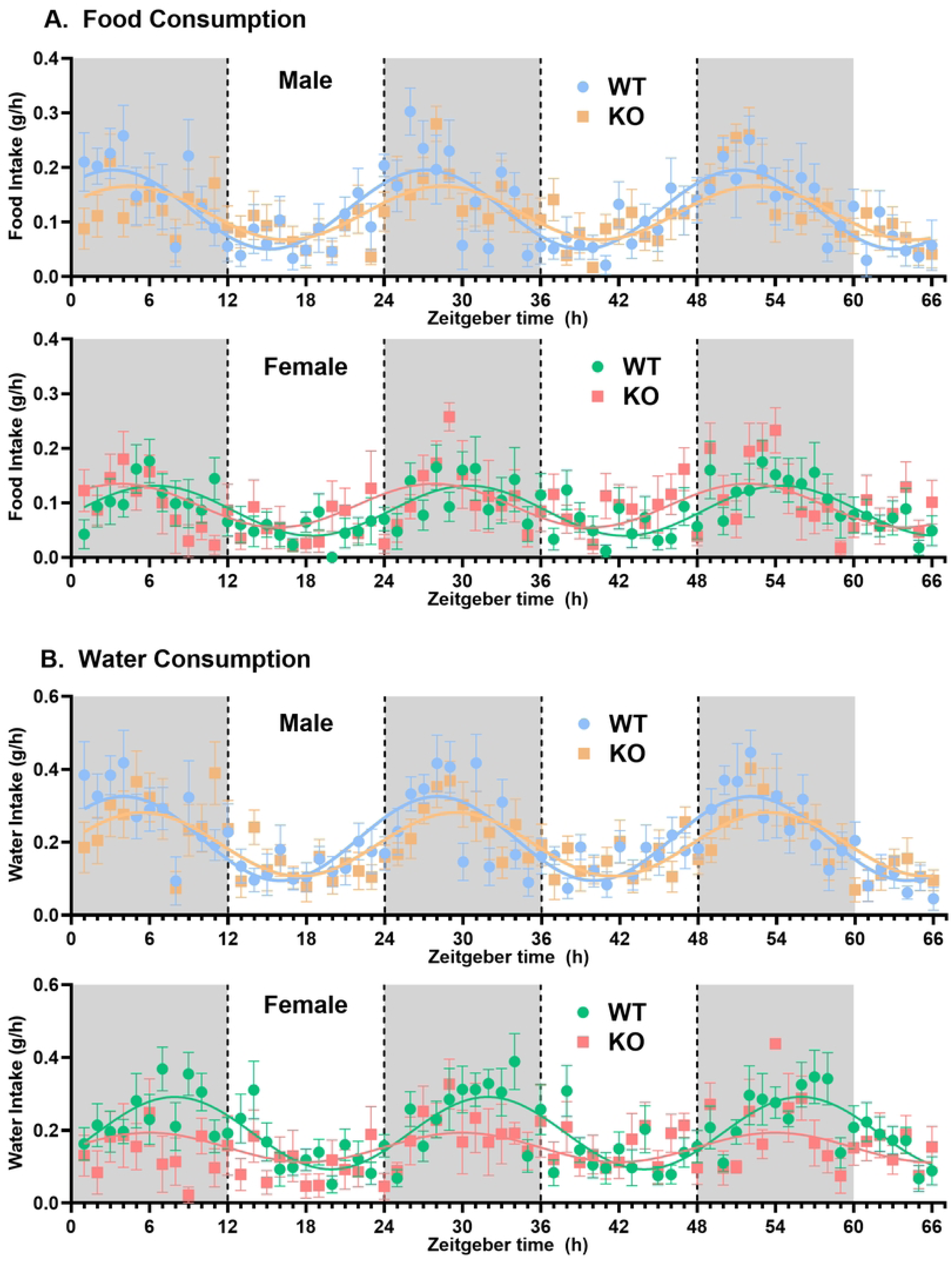
Effect of *slc43a3* deletion on food and water consumption in male and female WT and KO mice. Using an automated Columbus Lab Animal Monitoring System (CLAMS), the amount of food and water consumed were measured every 2-3 minutes over a period of 66 h (12 h L/D) and binned to obtain g/h. Zeitgeber time 0 was defined as the start of the first dark phase. Each point is the mean ± SEM from 8 mice. The lines represent LOWESS curve fits to the data using a 20-point smoothing window (GraphPad Prism v10.2). The area under the curve for these data sets are shown in Table 1.

**Table 1.**
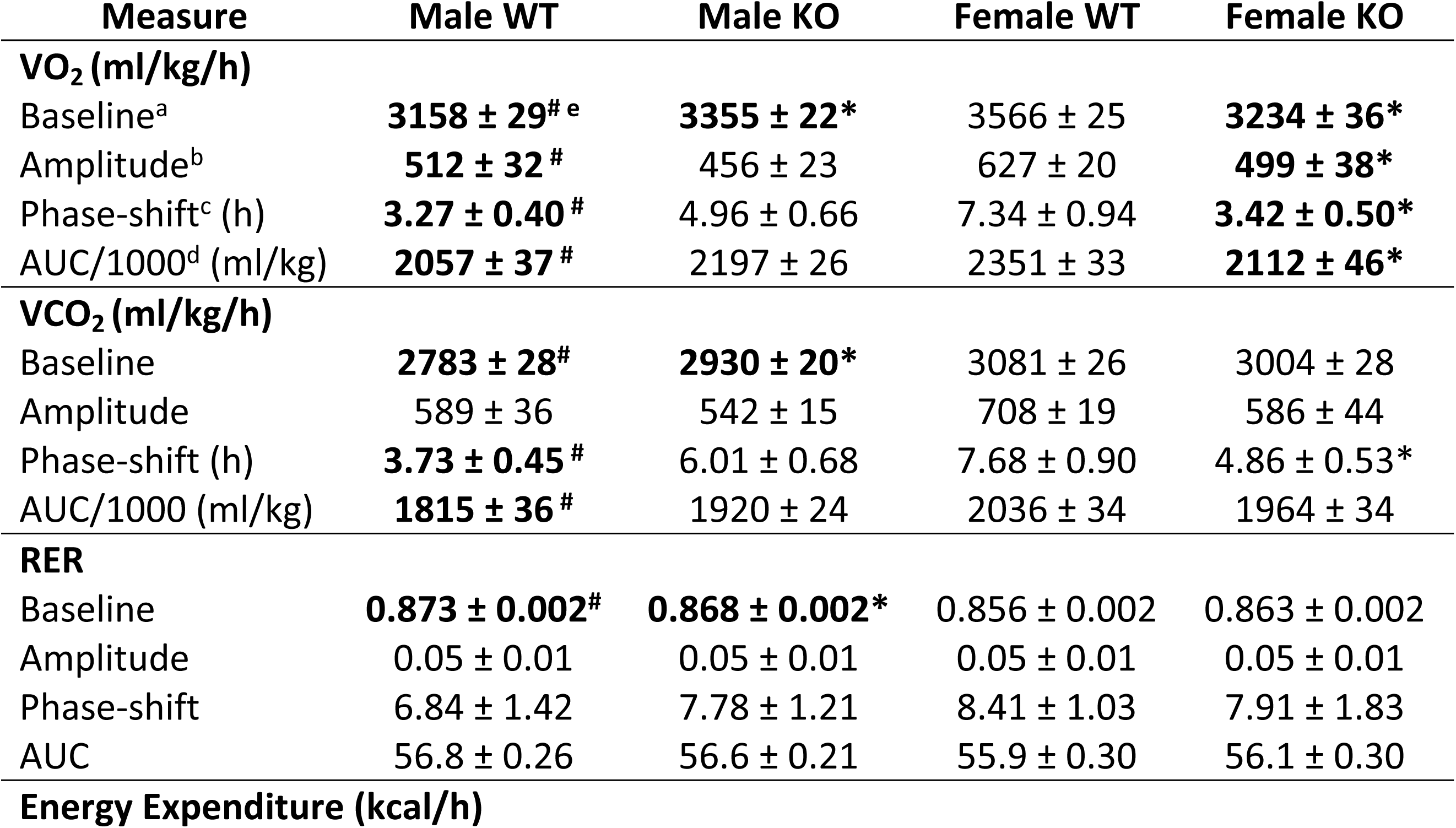
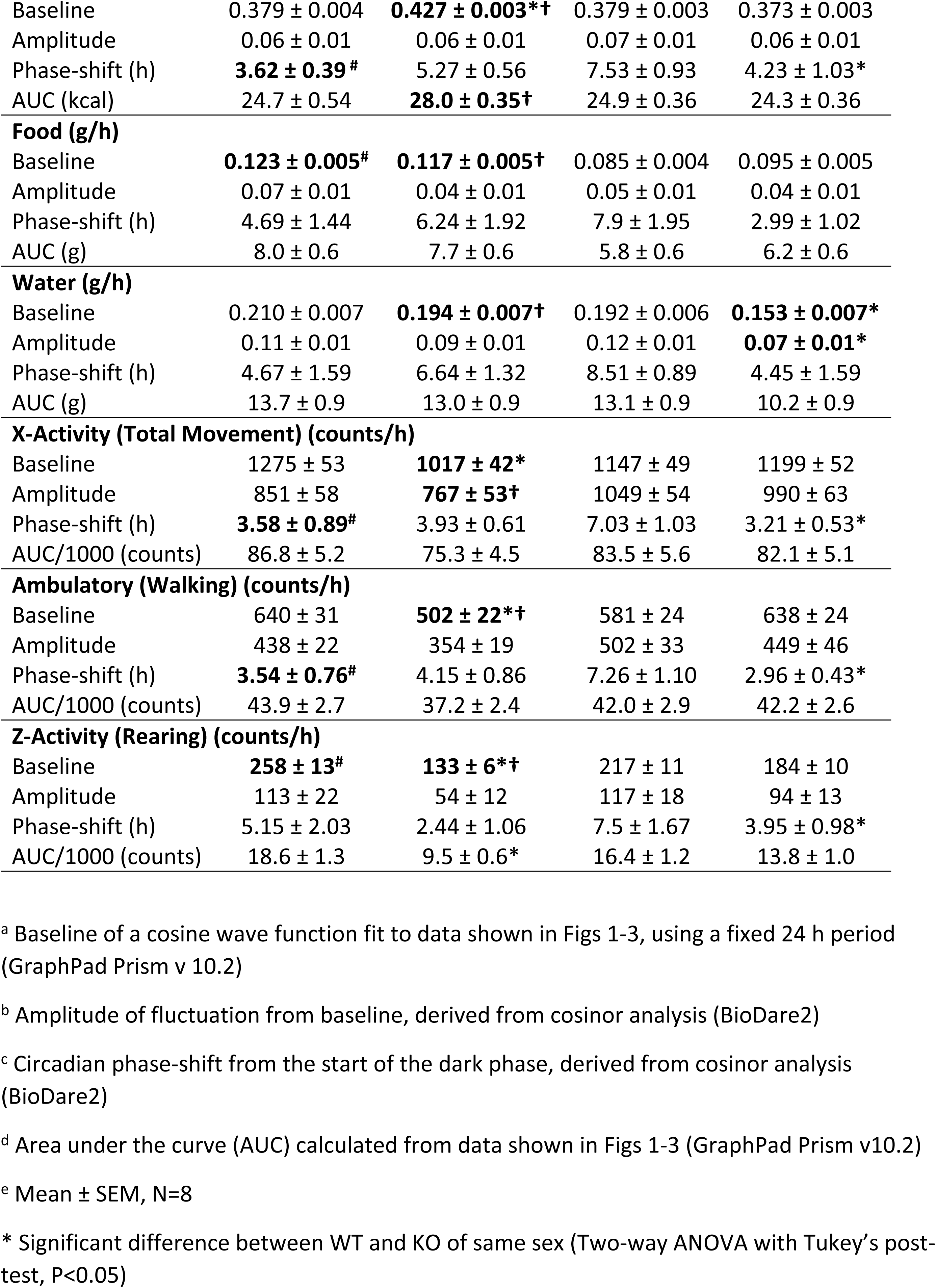
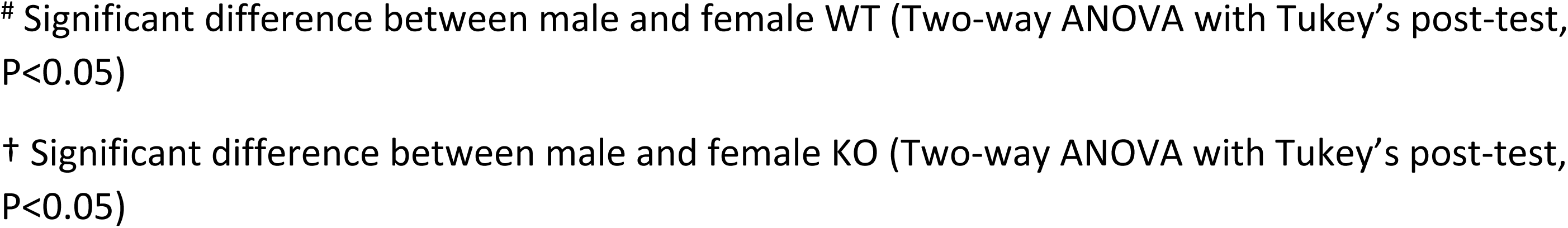
Summary of measured parameters.

| Measure | Male WT | Male KO | Female WT | Female KO |
| --- | --- | --- | --- | --- |
| **VO<sub>2</sub> (ml/kg/h)** |  |  |  |  |
| Baseline<sup>a</sup> | **3158 ± 29<sup>#e</sup>** | **3355 ± 22\*** | 3566 ± 25 | **3234 ± 36\*** |
| Amplitude<sup>b</sup> | **512 ± 32<sup>#</sup>** | 456 ± 23 | 627 ± 20 | **499 ± 38\*** |
| Phase-shift<sup>c</sup> (h) | **3.27 ± 0.40<sup>#</sup>** | 4.96 ± 0.66 | 7.34 ± 0.94 | **3.42 ± 0.50\*** |
| AUC/1000<sup>d</sup> (ml/kg) | **2057 ± 37<sup>#</sup>** | 2197 ± 26 | 2351 ± 33 | **2112 ± 46\*** |
| **VCO<sub>2</sub> (ml/kg/h)** |  |  |  |  |
| Baseline | **2783 ± 28<sup>#</sup>** | **2930 ± 20\*** | 3081 ± 26 | 3004 ± 28 |
| Amplitude | 589 ± 36 | 542 ± 15 | 708 ± 19 | 586 ± 44 |
| Phase-shift (h) | **3.73 ± 0.45<sup>#</sup>** | 6.01 ± 0.68 | 7.68 ± 0.90 | 4.86 ± 0.53* |
| AUC/1000 (ml/kg) | **1815 ± 36<sup>#</sup>** | 1920 ± 24 | 2036 ± 34 | 1964 ± 34 |
| **RER** |  |  |  |  |
| Baseline | **0.873 ± 0.002<sup>#</sup>** | **0.868 ± 0.002\*** | 0.856 ± 0.002 | 0.863 ± 0.002 |
| Amplitude | 0.05 ± 0.01 | 0.05 ± 0.01 | 0.05 ± 0.01 | 0.05 ± 0.01 |
| Phase-shift | 6.84 ± 1.42 | 7.78 ± 1.21 | 8.41 ± 1.03 | 7.91 ± 1.83 |
| AUC | 56.8 ± 0.26 | 56.6 ± 0.21 | 55.9 ± 0.30 | 56.1 ± 0.30 |
| **Energy Expenditure (kcal/h)** |  |  |  |  |
| Baseline | 0.379 ± 0.004 | **0.427 ± 0.003\*†** | 0.379 ± 0.003 | 0.373 ± 0.003 |
| Amplitude | 0.06 ± 0.01 | 0.06 ± 0.01 | 0.07 ± 0.01 | 0.06 ± 0.01 |
| Phase-shift (h) | **3.62 ± 0.39<sup>#</sup>** | 5.27 ± 0.56 | 7.53 ± 0.93 | 4.23 ± 1.03* |
| AUC (kcal) | 24.7 ± 0.54 | **28.0 ± 0.35†** | 24.9 ± 0.36 | 24.3 ± 0.36 |
| **Food (g/h)** |  |  |  |  |
| Baseline | **0.123 ± 0.005<sup>#</sup>** | **0.117 ± 0.005†** | 0.085 ± 0.004 | 0.095 ± 0.005 |
| Amplitude | 0.07 ± 0.01 | 0.04 ± 0.01 | 0.05 ± 0.01 | 0.04 ± 0.01 |
| Phase-shift (h) | 4.69 ± 1.44 | 6.24 ± 1.92 | 7.9 ± 1.95 | 2.99 ± 1.02 |
| AUC (g) | 8.0 ± 0.6 | 7.7 ± 0.6 | 5.8 ± 0.6 | 6.2 ± 0.6 |
| **Water (g/h)** |  |  |  |  |
| Baseline | 0.210 ± 0.007 | **0.194 ± 0.007†** | 0.192 ± 0.006 | **0.153 ± 0.007\*** |
| Amplitude | 0.11 ± 0.01 | 0.09 ± 0.01 | 0.12 ± 0.01 | **0.07 ± 0.01\*** |
| Phase-shift (h) | 4.67 ± 1.59 | 6.64 ± 1.32 | 8.51 ± 0.89 | 4.45 ± 1.59 |
| AUC (g) | 13.7 ± 0.9 | 13.0 ± 0.9 | 13.1 ± 0.9 | 10.2 ± 0.9 |
| **X-Activity (Total Movement) (counts/h)** |  |  |  |  |
| Baseline | 1275 ± 53 | **1017 ± 42\*** | 1147 ± 49 | 1199 ± 52 |
| Amplitude | 851 ± 58 | **767 ± 53†** | 1049 ± 54 | 990 ± 63 |
| Phase-shift (h) | **3.58 ± 0.89<sup>#</sup>** | 3.93 ± 0.61 | 7.03 ± 1.03 | 3.21 ± 0.53* |
| AUC/1000 (counts) | 86.8 ± 5.2 | 75.3 ± 4.5 | 83.5 ± 5.6 | 82.1 ± 5.1 |
| **Ambulatory (Walking) (counts/h)** |  |  |  |  |
| Baseline | 640 ± 31 | **502 ± 22\*†** | 581 ± 24 | 638 ± 24 |
| Amplitude | 438 ± 22 | 354 ± 19 | 502 ± 33 | 449 ± 46 |
| Phase-shift (h) | **3.54 ± 0.76<sup>#</sup>** | 4.15 ± 0.86 | 7.26 ± 1.10 | 2.96 ± 0.43* |
| AUC/1000 (counts) | 43.9 ± 2.7 | 37.2 ± 2.4 | 42.0 ± 2.9 | 42.2 ± 2.6 |
| **Z-Activity (Rearing) (counts/h)** |  |  |  |  |
| Baseline | **258 ± 13<sup>#</sup>** | **133 ± 6\*†** | 217 ± 11 | 184 ± 10 |
| Amplitude | 113 ± 22 | 54 ± 12 | 117 ± 18 | 94 ± 13 |
| Phase-shift (h) | 5.15 ± 2.03 | 2.44 ± 1.06 | 7.5 ± 1.67 | 3.95 ± 0.98* |
| AUC/1000 (counts) | 18.6 ± 1.3 | 9.5 ± 0.6* | 16.4 ± 1.2 | 13.8 ± 1.0 |

<sup>a</sup> Baseline of a cosine wave function fit to data shown in Figs 1-3, using a fixed 24 h period (GraphPad Prism v 10.2)
<sup>b</sup> Amplitude of fluctuation from baseline, derived from cosinor analysis (BioDare2)
<sup>c</sup> Circadian phase-shift from the start of the dark phase, derived from cosinor analysis (BioDare2)
<sup>d</sup> Area under the curve (AUC) calculated from data shown in Figs 1-3 (GraphPad Prism v10.2)
<sup>e</sup> Mean ± SEM, N=8
\* Significant difference between WT and KO of same sex (Two-way ANOVA with Tukey's post- test, P<0.05)

Another highly significant effect of deletion on *slc43a3* was a phase shift in the circadian rhythm of the female mice. This was apparent in the metabolic VO_2_, VCO_2_, and energy expenditure profiles, as well as in all of the activity measurements (Table 1, Figs 1, 2, 4, and 5); female WT and KO mice had circadian phase peaks of about 7.5 and 3.5 hours, respectively, after the start of the dark phase. This is also illustrated visually in Fig 4 for the VO_2_ data set, where it Is apparent that the loss of *slc43a3* eliminates the sex difference in the circadian phase. There was a similar trend in the food and water consumption in the female mice, but due to the greater variability in these data the change did not reach statistical significance (Fig 3, Table 1). A similar shift was not seen for the male KO versus WT mice. If anything, metabolic activity in the male KO mice tended to shift to a later circadian phase peak (∼5.5 hours) relative to male WT mice (∼3.5 hours) (Fig 5, Table 1). This shift in the circadian rhythm of the female KO mice eliminated the difference in circadian rhythm that was evident between the male and female WT mice (Fig 4). In general, male WT mice reached peak metabolic and physical activity about 3.5 hours into the dark phase, while female WT mice peaked at around 7.5 hours into the dark phase.

**Fig 4.**
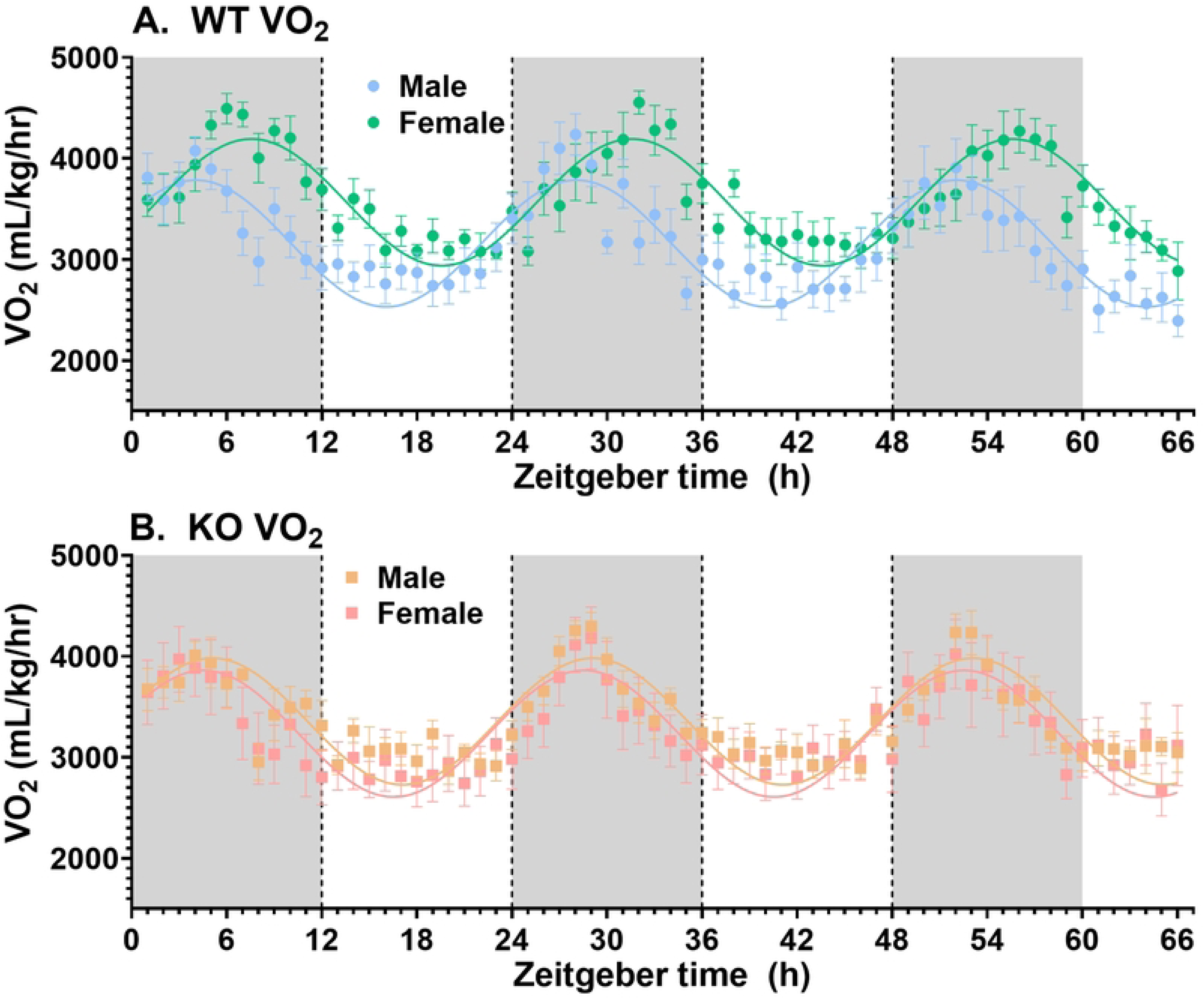
Effect of *slc43a3* deletion on the circadian rhythm of oxygen consumption (VO_2_) in male and female mice. Panel A shows the data from WT mice with a cosine wave function (fixed 24 h period) fitted to each data set. Panel B shows the data from KO mice with a cosine wave function fitted to each data set. Note that these data are a re-configuration of those shown in Fig 1A to illustrate the shift in circadian rhythm phase in the female mice upon deletion of *slc43a3*. Each point represents the mean ± SEM from 8 mice.

**Fig 5.**
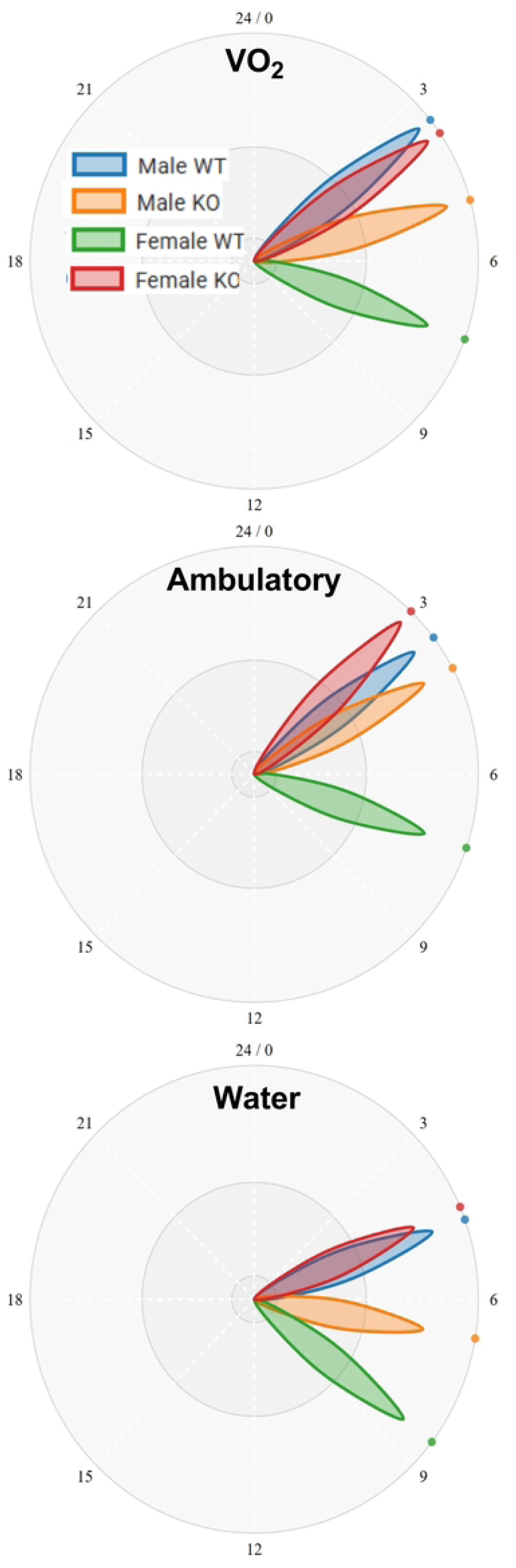
Circadian phase plots of the peak activity for VO_2_, ambulatory activity, and water consumption in male and female WT and KO mice. Plots were generated using BioDare2 based on cosinor analysis using the FFT NLLS method with linear detrending (biodare2.ed.ac.uk). Each phase ‘clock-hand’ represents the average time (h) after the start of the dark phase (time zero) derived from the data sets shown in Figures 1-3 and is the average from 8 mice. Mean ± SEM values for the phase-shifts are shown in Table 1.

Given the well-established role of purines in the regulation of cardiovascular function, we also compared the WT and KO mice with respect to their blood pressure and heart rate. Male KO mice showed a significant decrease (by about 20%) in both systolic and diastolic blood pressure, and a trend towards a decrease in heart rate relative to the male WT mice (Fig 6). In contrast, female WT and KO mice were not different with respect to their cardiovascular parameters.

**Fig. 6.**
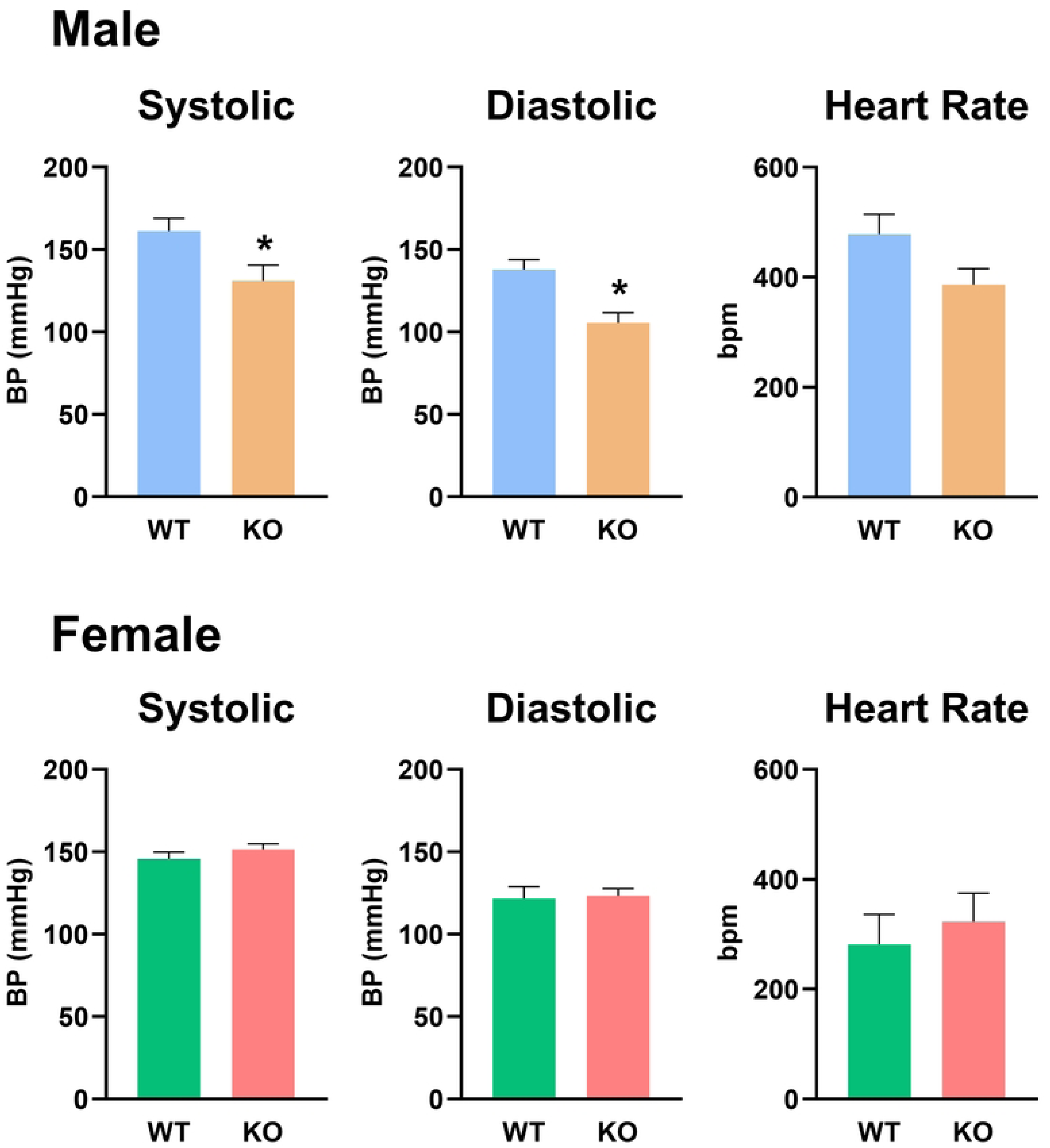
Effect of *slc43a3* deletion on resting blood pressure (systolic and diastolic) and heart rate of male and female WT and KO mice. Cardiovascular function was measured using a rodent tail cuff apparatus during the light-phase of the circadian cycle. * Indicates a significant difference between WT and KO mice (One-way ANOVA with Tukey’s post-test, P<0.05).

## Discussion

The metabolic and behavioural characteristics of the mice used in this study were consistent with previously published data on young adult C57BL/6J mice [9], including the approximate 4 hr circadian phase angle difference between male and female mice seen for many of the parameters measured [10]. While no gross abnormalities were observed in the KO mice, analyses of metabolic function and activity profiles showed significant impacts of the loss of *slc43a3*. There were also noteworthy differences between WT male and female mice with respect to a number of the parameters assessed. The most dramatic effect of *slc43a3* deletion was the loss of the sex difference in metabolic function seen for the WT mice. Female WT mice had consistently higher VO_2_ and VCO_2_ values and a higher degree of fluctuation (amplitude from cosine wave analysis) of these parameters over the 24-hr diurnal cycle. In contrast, female KO mice were not significantly different from male WT or KO mice in this regard. Male mice were not similarly impacted by the loss of *slc43a3*. *Slc43a3* encodes for an equilibrative nucleobase transporter that is involved in the salvage of purine nucleobases (e.g. adenine) by cells to support cellular metabolic activity. Thus, it is not surprising that its loss would have an attenuating impact on oxygen consumption (VO_2_) and carbon dioxide generation (VCO_2_) which reflect overall cellular metabolic activity in the mice. The sex specific nature of this effect is intriguing though. We have previously shown that male and female WT mice have similar levels of expression of *slc43a3*, and that the loss of this gene does not impact the expression of adenine phosphoribosyltransferase (APRT), the critical enzyme responsible for catabolism of adenine to AMP [4], in either sex [6]. This suggests that female mice, in this case, may be more reliant on nucleobase salvage (as opposed to *de novo* purine synthesis) to maintain their higher metabolic activity.

The circadian phase angle difference between male and female WT mice was also lost in the KO mice for all of the parameters measured. The circadian rhythm in rodents is controlled by the hypothalamic suprachiasmatic nucleus [11]. While the molecular/cellular signalling pathways involved in the control of circadian rhythms are complex [12], there are several mechanisms by which disruption of the purine metabolic pathways (via reduced salvage of purine nucleobases by ENBT1) may impact circadian rhythm. This is particularly important considering that adenine is a metabolic precursor to adenine nucleotides. Indeed adenosine-receptor mediated signaling itself has been implicated in the regulation of the circadian clock [13]. Adenosine 3’,5’-monophosphate kinase (AMPK) phosphorylates the transcription factors PER and CRY that are integral to circadian clock control, thus promoting their polyubiquitination and subsequent degradation[14, 15]. AMPK activity is regulated by the ratio of AMP:ATP in cells [16], which is sensitive to purine nucleoside/nucleobase levels [17]. Furthermore, calcium/cAMP response elements have been found in the promoters of several clock genes, and cAMP-dependent signaling has been proposed as a core component of the mammalian circadian pacemaker[18]. Adenosine can regulate intracellular calcium levels [19, 20] and is a precursor to cAMP. Our finding that it is specifically the female circadian rhythm that is phase-shifted in the *slc43a3-null* mice, implies that estrogen plays a role in this. While sex differences in circadian rhythms have been well documented [21, 22], there is notably less information in the literature regarding the underlying mechanisms for these differences. While still speculative, a possible linkage between purine metabolism, estrogen, and circadian rhythm may involve AMPK. The observed disruption of metabolic activity in the female mice upon loss of *slc43a3* may impact AMPK activity in the female mice specifically leading to changes in circadian rhythm control. It is also noteworthy that AMPK has been shown to be regulated by estrogen [23], which may also be a factor in the sex-specific differences noted in the present study.

The other sex-specific difference observed, was the significantly decreased rearing activity seen in the male *slc43a3* KO mice, and a trend towards decreased activity in general in this group of mice. This was accompanied by a significant decrease in both systolic and diastolic blood pressure in the male KO mice, and an increase in overall energy expenditure. While interpretation is confounded somewhat by the fact that blood pressure was measured during the light phase and rearing activity occurred primarily during the dark phase, it is tempting to speculate that the decrease in blood pressure may result in a postural hypotension type of response in these mice, reducing their preference for elevating their bodies. Alternatively, it has been reported that chronic low blood pressure in mice reduces their performance in rotarod tests[24], suggesting problems with maintaining balance which might also decrease the frequency/duration of rearing. This requires further investigation using free-roaming telemetry monitoring of blood pressure and positional information.

While mechanistic underpinnings of the changes observed still require further investigation, the loss of *slc43a3* in mice clearly results in significant changes in metabolic activity in female mice and changes in cardiovascular function and physical activity measures in male mice. These data highlight the importance of the purine nucleobase transporter ENBT1 in the endogenous regulation of metabolic function.

## Acknowledgments

The technical assistance of Tierah Hinchliffe and Hannah Dean in support of this work is gratefully acknowledged. Much of the work presented in this study was conducted in the Metabolic Core Facility of the Cardiovascular Research Centre, University of Alberta, with the guidance of facility technologist Amy Barr.

